# mTOR inhibitors as radiosensitizers in neuroendocrine neoplasms

**DOI:** 10.1101/2020.07.01.181727

**Authors:** Samantha Exner, Gerard Arrey, Vikas Prasad, Carsten Grötzinger

**Affiliations:** Department of Hepatology and Gastroenterology, Charité – Universitätsmedizin Berlin, Berlin, Germany; Department of Nuclear Medicine, Charité – Universitätsmedizin Berlin, Berlin, Germany; Department of Nuclear Medicine, University Hospital Ulm, Ulm, Germany; Molecular Cancer Research Center (MKFZ), Charité – Universitätsmedizin Berlin, Berlin, Germany; German Cancer Consortium (DKTK), Berlin, Germany; German Cancer Research Center (DKFZ), Heidelberg, Germany

**Keywords:** Neuroendocrine neoplasms, mTOR inhibitors, PRRT, radiosensitizer

## Abstract

Peptide receptor radioligand therapy (PRRT) has evolved as an important second-line treatment option in the management of inoperable and metastatic neuroendocrine neoplasms (NEN). Though high radiation doses can be delivered to the tumors, complete remission is still rare. Radiosensitization prior to PRRT is therefore considered to be a promising strategy to improve the treatment effect. In this study, effect and mechanism of mTOR inhibitors were investigated in a comprehensive panel of five NEN cell lines (BON, QGP-1, LCC-18, H727, UMC-11), employing assays for cellular proliferation, clonogenic survival, cell cycle modification and signaling. mTOR inhibition lead to growth arrest with a biphasic concentration-response pattern: a partial response at approximately 1 nM and full response at micromolar concentrations (8-48 μM). All cell lines demonstrated elevated p70S6K phosphorylation yet also increased phosphorylation of counterregulatory Akt. The pulmonary NEN cell line UMC-11 showed the lowest induction of phospho-Akt and strongest growth arrest by mTOR inhibitors. Radiation sensitivity of the cells (50% reduction versus control) was found to range between 4 and 8 Gy. Further, mTOR inhibition was employed together with irradiation to evaluate radiosensitizing effects of this combination treatment. mTOR inhibition was found to radiosensitize all five NEN cells in an additive manner with a moderate overall effect. The radiation-induced G2/M arrest was diminished under combination treatment, leading to an increased G1 arrest. Further investigation involving a suitable animal model as well as radioligand application such as ^177^Lu-DOTATATE or ^177^Lu-DOTATOC will have to demonstrate the full potential of this strategy for radiosensitization in NEN.

## Introduction

Peptide receptor radioligand therapy (PRRT) has evolved as an important second-line treatment option in the management of inoperable and metastatic neuroendocrine neoplasms (NEN). In one of the earliest studies, comprising 310 GEP-NET patients, treatment with ^177^Lu-DOTATATE resulted in complete tumor remission in 2 % and partial tumor remission in 28 % of the patients, with a median progression-free survival of 40 months. Side effects were found to be mild (1). Recently, the first randomized multicenter PRRT trial (NETTER-1) evaluated the effect of combined somatostatin analogs (SSA) and PRRT treatment in comparison to SSA alone in patients with advanced, progressive midgut NENs. This phase III trial confirmed previously obtained results of various studies, demonstrating high response rates and increased progression-free survival after PRRT (2). Though high radiation doses (up to 250 Gy) can be delivered to the tumors, complete remission is still rare (3). Radiosensitization prior to PRRT is therefore considered to be a promising strategy to improve the treatment effect.

The serine/threonine kinase mTOR (mammalian target of rapamycin) is a central integrator of environmental signals such as nutrients, growth factors and stress, coordinating cell growth and proliferation. Depending on the proteins it is interacting with, it can form the two complexes mTORC1 and mTORC2 which have their own distinct downstream effectors. Activated mTORC1 initiates protein translation by phosphorylation of S6K1 (also known as p70S6K) and 4EBP1, which in turn further engage S6 ribosomal protein and eIF4E. At the same time, activated S6K1 negatively regulates the PI3K-Akt pathway by inhibition of IRS. Furthermore, mTORC1 promotes cell cycle progression, inhibits autophagy and controls transcription and the DNA damage response. On the other hand, mTORC2 is in charge of cell survival, metabolism and the actin cytoskeleton (4,5).

Dysregulated mTOR signaling has been demonstrated in various cancers, which made mTOR a promising target and facilitated the development of derivatives of rapamycin, a naturally occurring mTOR inhibitor. Two examples for FDA-approved rapalogs with improved pharmacological and solubility qualities are temsirolimus (CCI-779) and everolimus (RAD001) (6). They form a complex with FKBP12 before binding to mTORC1. Inhibition of mTORC1 leads to G1 cell cycle arrest, reduced tumor angiogenesis, apoptosis induction and enhanced sensitivity towards DNA-damaging agents (4). However, rapalogs interrupt not only downstream functions, but also the S6K1 feedback loop. This results in an upregulation of Akt-mediated pro-survival signaling and may counteract the antitumor activity of the inhibitor (7). While rapalogs act as universal inhibitors of mTORC1, Akt downregulation was observed only in a few cancer cell lines, indicating a cell-type specific inhibition of mTORC2 (8). Possibly, cells with PTEN loss and hyperactive PI3K-Akt signaling may be more dependent on mTORC1 and show higher sensitivity towards rapalogs (9). Also, other signaling pathways such as the MAPK cascade seem to be activated by mTOR inhibition, although this is less well investigated (10).

Mutations as well as aberrant activations in the PI3K-Akt-mTOR network were also observed in NENs (11). A series of clinical trials (RADIANT) resulted in the FDA-approval of everolimus for advanced pancreatic, non-functional gastrointestinal and lung NETs (12–15). Treatment with everolimus prolonged median progression-free survival by 6.4 to 7.1 months when compared to the placebo group. Response rates of temsirolimus were similar to those observed with everolimus, as evaluated in a phase II study in advanced NENs (16). However, the authors concluded that temsirolimus, when applied as a single drug, may yield only modest clinical benefit (17). In addition, it has to be administered intravenously, whereas everolimus is available as an oral formulation (18,19). Recently, a small phase I study assessed the safety and optimal dose for a combined treatment of NENs with everolimus and PRRT (^177^Lu-DOTATATE) in 16 patients. Overall response was observed in 44 % of patients, and the maximum tolerated dose for everolimus in combination was found to be 7.5 mg daily (20).

In this study, effect and mechanism of mTOR inhibitors were investigated in a comprehensive panel of five NEN cell lines, employing assays for cellular proliferation, clonogenic survival, cell cycle modification and signaling. Further, mTOR inhibition was employed together with irradiation to evaluate radiosensitizing effects of this combination treatment.

## Material and Methods

### Antibodies

| Designation | Species | Dilution | Cat No/Manufacturer |
| --- | --- | --- | --- |
| *Primary antibodies* |  |  |  |
| Akt | Rabbit, polyclonal | 1:1000 in 5% BSA/TBST | #9272, Cell Signaling (Danvers, US) |
| Caspase-3 | Rabbit, polyclonal | 1:1000 in 5% milk/TBST | #9662, Cell Signaling (Danvers, US) |
| P44/42 MAPK (ERK1/2) (3A7) | Mouse, monoclonal | 1:2000 in 5% milk/TBST | #9107, Cell Signaling (Danvers, US) |
| p70 S6 Kinase | Rabbit, polyclonal | 1:1000 in 5% BSA/TBST | #9202, Cell Signaling (Danvers, US) |
| PARP | Rabbit, polyclonal | 1:1000 in 5% milk/TBST | #9542, Cell Signaling (Danvers, US) |
| Phospho-Akt (Ser473) (D9W9U) | Mouse, monoclonal | 1:1000 in 5% BSA/TBST | #12694, Cell Signaling (Danvers, US) |
| Phospho-p44/42 MAPK (ERK1/2) (Thr202/Tyr204) (D13.14.4E) XP | Rabbit, monoclonal | 1:2000 in 5% BSA/TBST | #4370, Cell Signaling (Danvers, US) |
| Phospho-p70 S6 Kinase (Thr389) (1A5) | Mouse, monoclonal | 1:1000 in 5% milk/TBST | #9206, Cell Signaling (Danvers, US) |
| *Secondary antibodies* |  |  |  |
| Anti-rabbit IgG (H+L) horseradish peroxidase conjugated | Goat, polyclonal | 1:5000 in 5% milk/TBST | #111035003 Jackson ImmunoResearch (West Grove, US) |
| Anti-mouse IgG (H+L) horseradish peroxidase conjugated | Goat, polyclonal | 1:2000 in 5% milk/TBST | #115035003 Jackson ImmunoResearch (West Grove, US) |

### Protein quantification

Protein concentration was determined using the BCA Protein Assay Kit (Pierce ThermoFisher, Waltham, MA, USA) according to the manufacturer’s protocol. Colorimetric changes were measured at 562 nm and the obtained OD values were used to interpolate protein concentrations from a standard curve prepared with BSA (0-10 mg/ml).

### SDS-PAGE

Protein gel electrophoresis was performed according to standard protocols. Cell lysates were separated on discontinuous SDS polyacrylamide gels. 10 μg of protein were diluted in 5x sample buffer and water in 10 μl total volume, denatured for 5 min at 95 °C and spun down shortly. Polymerized gels were assembled in a Mini-PROTEAN Tetra tank (Bio-Rad, Hercules, CA, USA), the upper chamber filled with top buffer (0.1 M Tris-HCl, 0.1 M Tricine pH 8.25, 0.1 % v/w SDS) and the samples loaded into the lanes. The lower chamber was filled with bottom buffer (0.2 M Tris-HCl, pH 8.8) and the gels run at 100 V. The Precision Plus Kaleidoscope Standard (Bio-Rad) was used as molecular weight marker.

### Western blotting

Electrophoretically separated proteins were transferred to nitrocellulose membranes by the wet blot technique. For this, polyacrylamide gel, membrane, fiber pads and filter paper were equilibrated in transfer buffer (10 % v/v Rotiphorese 10x SDS-PAGE, 20 % 96% ethanol in water) for 15 min, stacked and locked into a cassette. The cassette was placed into a mini Trans-Blot tank together with an ice pad and a stir bar and filled with transfer buffer. Blotting was performed for 1 h at 100 V and constant 350 mA on a magnetic stirrer. Membrane was stained with ponceau S for assessment of homogenous protein transfer, destained with water and incubated in blocking buffer (5 % w/v nonfat dry milk in TBST) for 60 min. After a short wash with TBST, membrane was incubated with primary antibody overnight at 4 °C. Membrane was washed three times for 5 min with TBST, incubated with HRP-coupled secondary antibody diluted in blocking buffer for 60 min at room temperature and again, washed three times for 5 min with TBST. Chemiluminescence was induced by addition of 500 μl SuperSignal West Dura (Pierce ThermoFisher) substrate and captured with a VersaDoc imaging system (Bio-Rad). Protein signals were analyzed and quantified with Image Lab software (Bio-Rad).

### Drug and radiation treatment

Cells were seeded at distinct densities in 96-, 12- or 6-well plates, depending on the experimental setup, and grown overnight. Substances were added in medium on top of the wells in double concentration for the indicated final concentrations. Irradiation was performed using an external caesium-137 source at a dose rate of 1 Gy/min and further incubation without medium change. For combination treatments, cells were incubated for 24 h with the respective substance before radiation or radioligand treatment was applied.

### Cell cycle analysis

Cell cycle analysis was performed on the basis of DNA content measurement by flow cytometry. The fluorescence intensity of DNA binding dyes such as propidium iodide is proportional to the DNA content of the cell, thereby allowing the discrimination of cells in sub-G1, G1, S and G2/M cell cycle phases (21). Cells were seeded in 12-well plates at a density between 100,000 to 200,000 cells per well and grown overnight before treatment. Both supernatant and cells were harvested at distinct time points and fixated with ice cold 70 % ethanol, which was added dropwise while vortexing to avoid cell aggregation. After fixation at −20 °C for at least 24 h, samples were washed with PBS and stained with propidium iodide solution (20 μg/ml propidium iodide, 20 μg/ml RNase A in PBS), containing RNaseA to remove interfering RNA. For each sample 10,000 events were counted with a flow cytometer measuring forward and sideward scatter as well as integrated (area, FL2-A) and pulse (width, FL2-W) red fluorescence. Doublet discrimination was performed by gating the cells using FL2-A vs. FL2-W, in that way excluding two aggregating G1 cells that appear to be one single G2/M cell (22). The gate was applied to the PI histogram, cell cycle phases marked and the percentages of cells in each phase quantified with CellQuest Pro software (Becton Dickinson).

### Cell viability – metabolic activity and cell number

Cells were seeded in quadruplicates at a density of 5,000 cells in 50 μl medium per well in 96-well plates, grown overnight and treated as described. After 96 h, metabolic activity was determined by addition of 100 μl medium containing 0.4 mM of the redox indicator resazurin. Cells were incubated for 3-4 hours and the resulting fluorescence was measured with an EnVision Multilabel Plate Reader (excitation filter: TRF 495 nm, emission filter: dysprosium 572 nm). Subsequently, the supernatant was removed, cells were fixated with 4 % formaldehyde for 10 min, stained with DAPI (1 μg/ml in PBS/0.1 % Triton) for another 10 min and wells were covered with 80 μl PBS for image acquisition. Four fields per well were imaged in an IN Cell Analyzer 1000 (GE, Reading, UK) using the 4x objective. Image stacks were analyzed and nuclei counted by IN Cell software. The values were averaged and normalized as percent of control treated with vehicle.

### Clonogenic survival

In comparison to short term cell viability assays, the clonogenic survival assay evaluates the ability of single cells to reproduce and form colonies, so called clones. Cells were seeded in duplicates and a density of 5,000 cells in 500 μl medium per well in 12-well plates and treated as described. Cells were incubated without medium change for 1-2 weeks. Finally, colonies were fixed with 70 % ethanol for 10 min, stained with crystal violet solution solution (0.2 % w/v crystal violet, 20 % methanol in water) for another 10 min and carefully rinsed with tap water. Plates were dried overnight and digitized with an Odyssey infrared scanner (700 nm channel, intensity 3, 84 μm resolution and medium quality). For quantification, images were analyzed using the ColonyArea plugin for ImageJ (23).

### Statistics and data availability

If not indicated otherwise, dose-response curves were plotted with GraphPad Prism 5.3 and the data fitted using nonlinear regression and the variable slope model (four-parameters). As x values, base 10 logarithms of doses or concentrations were entered. Statistical analyses were performed with the same software. The quantitative data for this study have been permanently published in a public repository accessible via this link: https://doi.org/10.5281/zenodo.3922212

## Results

### Effect of mTOR inhibitors on NEN cells

To evaluate the effect of the mTOR inhibitors temsirolimus and everolimus on neuroendocrine tumor cells, five NEN cell lines from different organs of origin were studied: BON and QGP-1 (both from pancreas), LCC-18 (from colon), and H727 and UMC-11 (both from lung). The cells were incubated with either mTOR inhibitor and two parameters of cell viability were determined 96 hours after the start of the incubation: metabolic activity and cell number. In both assays, temsirolimus and everolimus led to a biphasic inhibition of cell viability in all five NEN cell lines (**Figure 1**), displaying similar concentration-response curves. Metabolic activity as well as cell number decreased while inhibitor concentrations increased, with two calculated IC_50_ values in the nanomolar and micromolar range, respectively (Table 1). The low nanomolar IC_50_ differed only slightly between cell lines and assays (around 1 nM), whereas the high micromolar IC_50_ demonstrated greater variation. Here, values ranged from 8 to 21 μM for temsirolimus and from 30 to 48 μM for everolimus. At nanomolar concentration, the inhibitors reduced cell viability by 20-75 %, with BON being the most resistant cell line (20 %) and UMC-11 the most sensitive (75 %). In contrast, when applying micromolar concentrations, all NEN cell lines eventually showed a complete loss of cell viability.

**Figure 1:**
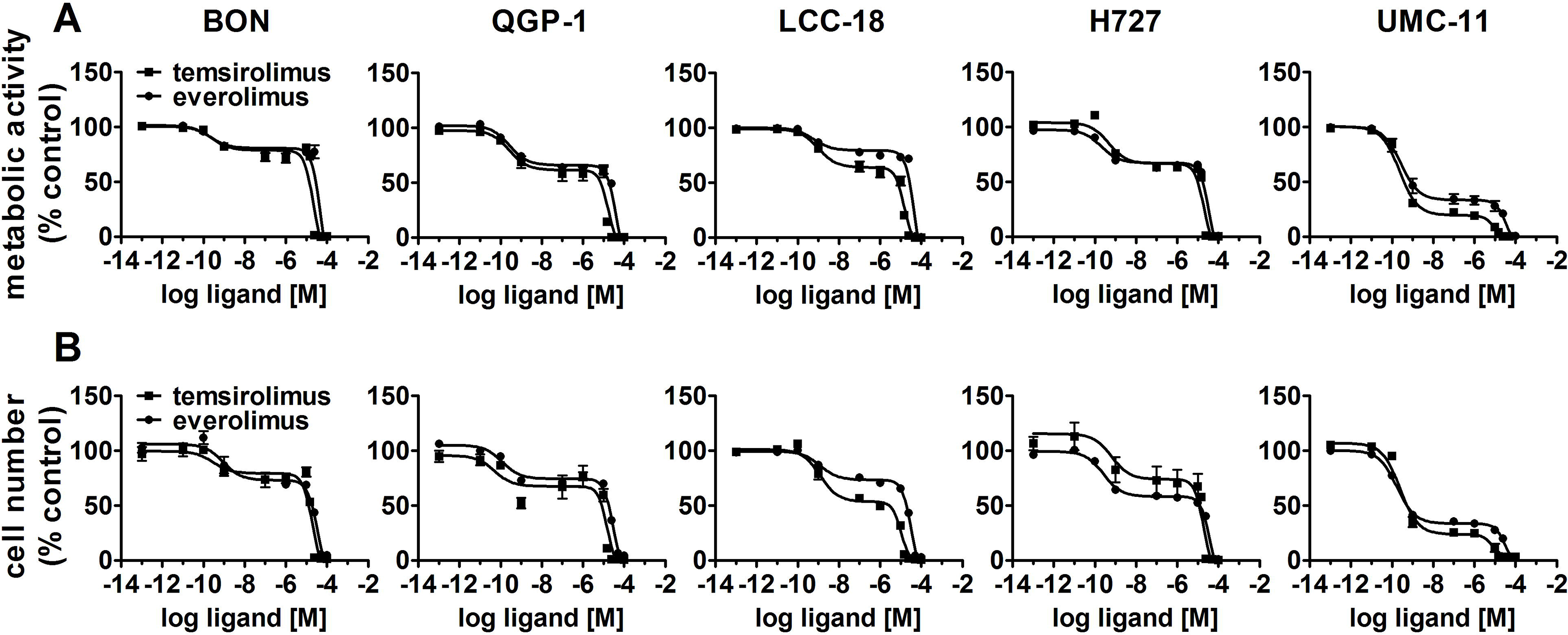
Treatment with mTOR inhibitors results in a biphasic inhibition of NEN cell viability. NEN cell lines were treated with increasing concentrations of temsirolimus or everolimus (0.1 pM to 100 μM), incubated for 96 h and analyzed for metabolic activity **(A)** and cell number **(B)**. Data represent mean ± S.E.M. (n=3).

**Table 1.**
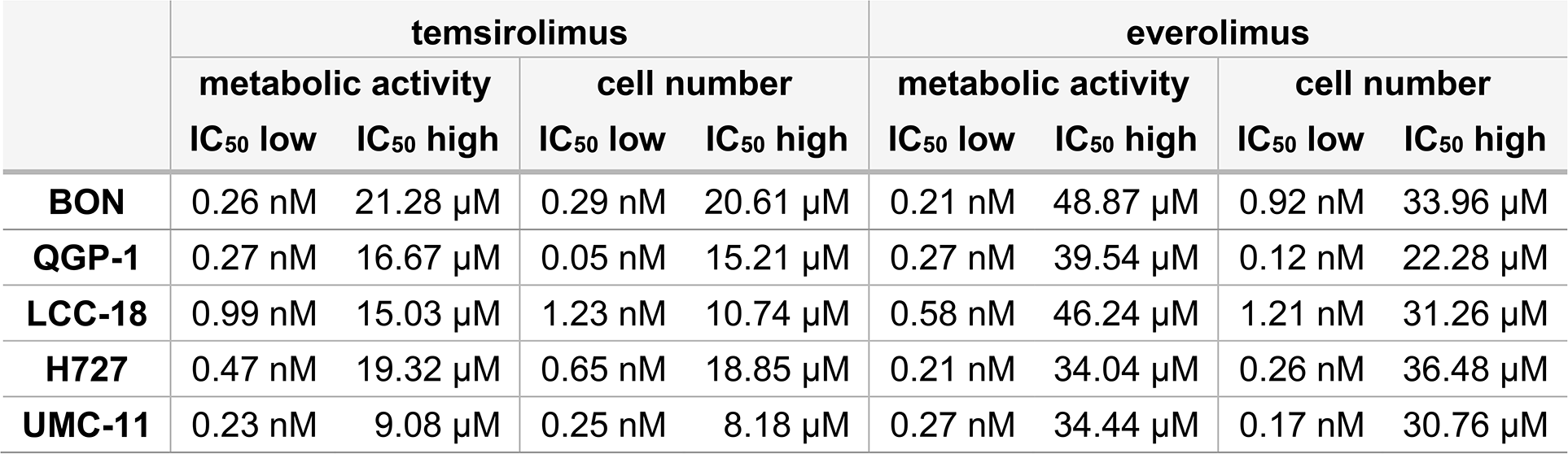
Summary of IC_50_ values for mTOR inhibitors.

To investigate the inhibitors’ impact on long-term cell survival and proliferation, clonogenic assays were performed. Both inhibitors led to similar concentration-response curves, with the most profound effect on UMC-11 and H727 cells (Figure 2). In contrast to the cell viability assays, even low nanomolar concentrations (1 nM, 10 nM) strongly inhibited cell proliferation in these pulmonary NEN cell lines; by 40-70 % for H727 and even by 85-95 % for UMC-11. When applying micromolar concentrations, no colonies were detectable at all. The other cell lines proved to be more resistant to treatment with nanomolar concentrations, but showed moderate inhibition of 60 % to 70 % in the micromolar range.

**Figure 2:**
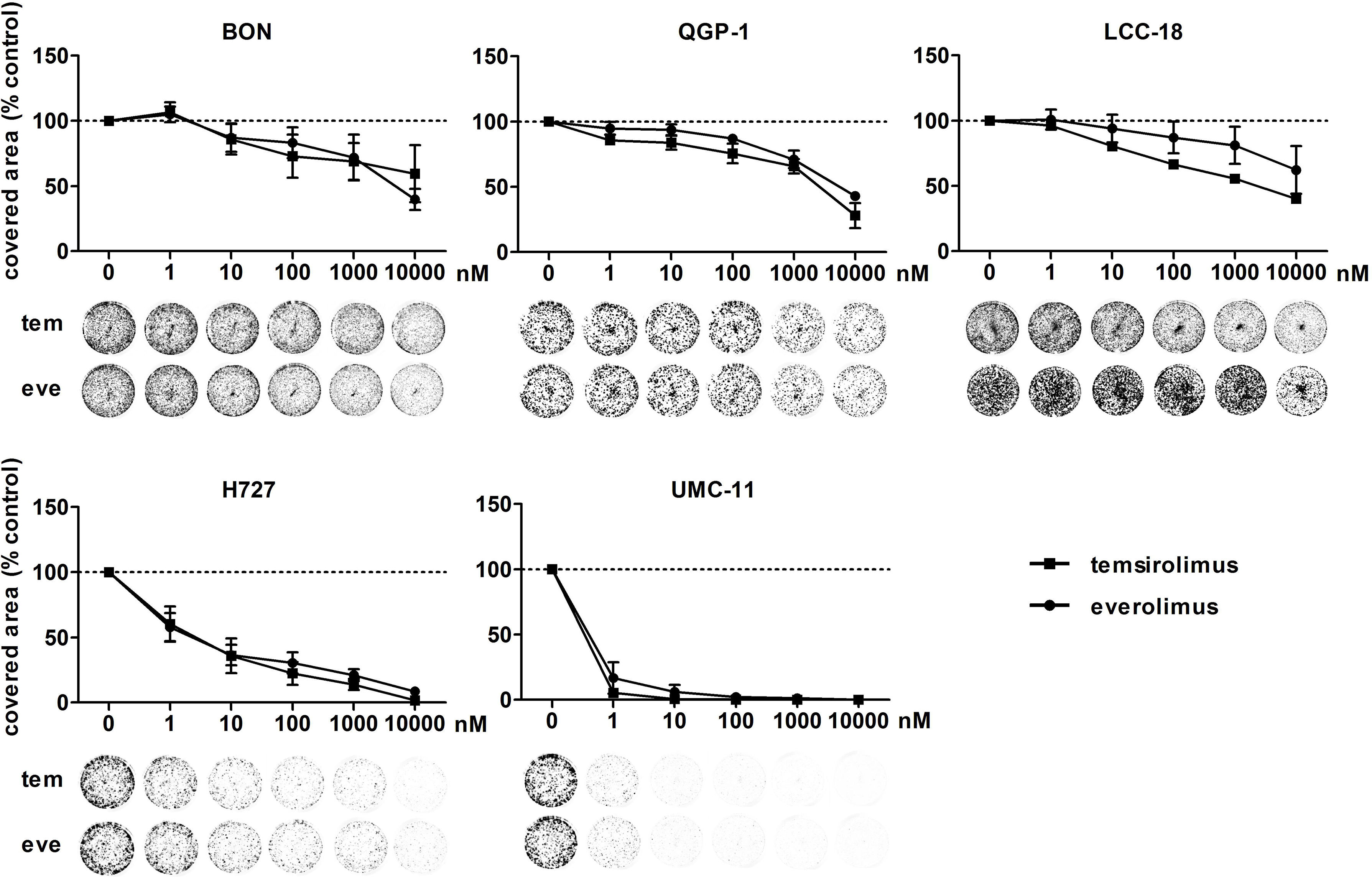
Clonogenic survival of NEN cell lines is affected by mTOR inhibitors. NEN cell lines were seeded at low density, treated with increasing concentrations of temsirolimus (tem) or everolimus (eve) and incubated for 1-2 weeks until colony formation. Data were normalized to untreated controls and represent mean ± S.E.M. (n=3-4) or mean only (LCC-18 treated with temsirolimus, n=1).

As mTOR inhibitors are known to inhibit cell cycle progression in the G1 phase, their effect in the NEN cell lines was investigated. Cell cycle analysis by flow cytometry was performed 24 h to 96 h after treatment. Within 24 hours, temsirolimus induced a moderate increase of cells in the G1 phase (Figure 3, Table 2). However, this turned out to be a transient effect as percentages leveled out after 72 h, at the latest. H727 cells were not affected at all, with the G1 fraction remaining almost the same during the whole period.

**Figure 3:**
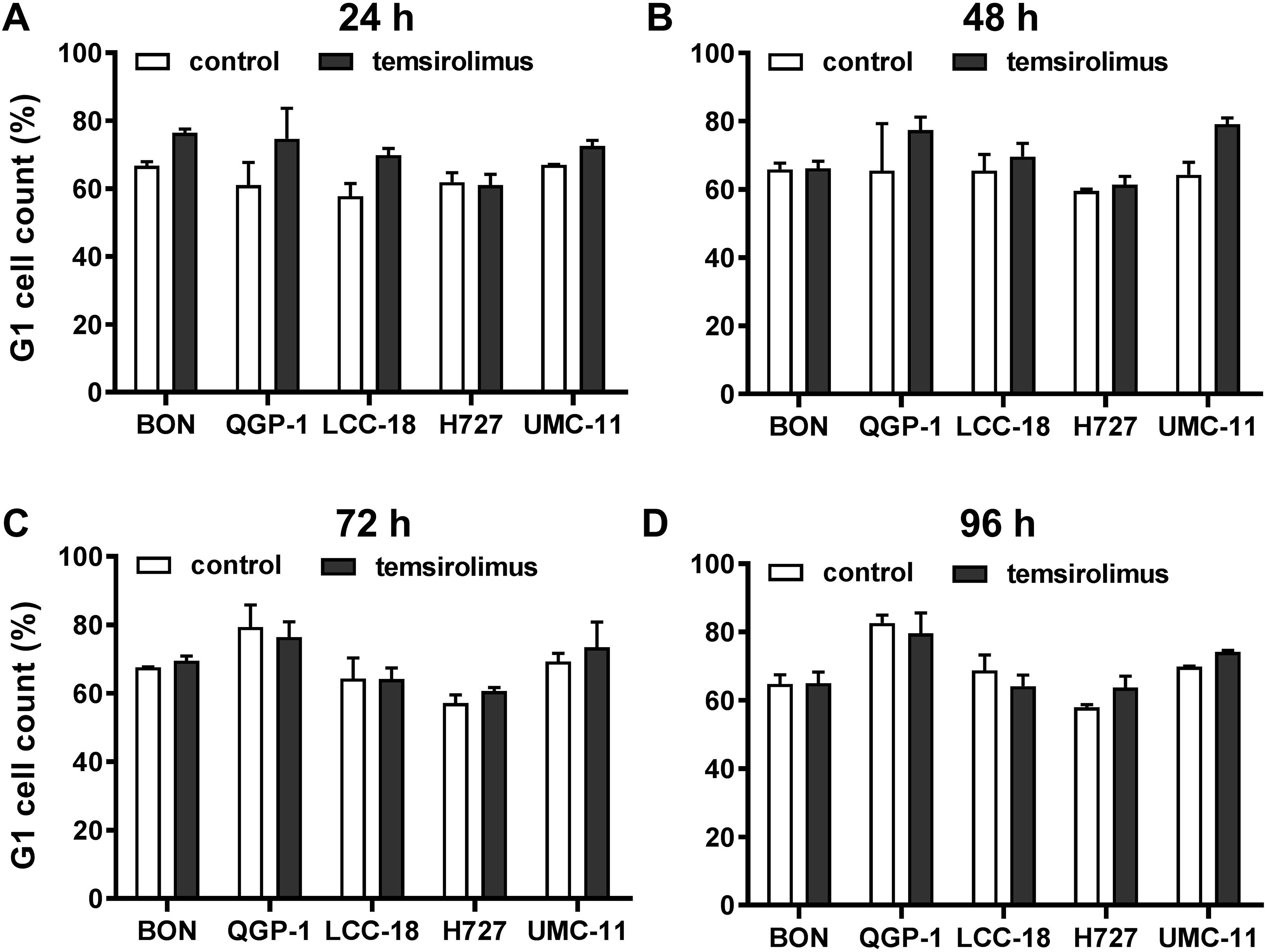
Increased accumulation of cells in G1 after mTOR inhibitor treatment. Five NEN cell lines were treated with vehicle (control) or 1 μM temsirolimus. After **(A)** 24, **(B)** 48, **(C)** 72 and **(D)** 96 h, incubation samples were collected for cell cycle analysis by flow cytometry. Data show mean ± S.E.M. of n=2-3.

**Table 2.** Percentages of cells in G1 cell cycle after mTOR inhibitor treatment. Data show mean ± S.E.M. (n=2-3).

|  | 24h |  | 48h |  | 72h |  | 96h |  |
| --- | --- | --- | --- | --- | --- | --- | --- | --- |
|  | 0 | 1 µM | 0 | 1 µM | 0 | 1 µM | 0 | 1 µM |
| **BON** | 66.8 ± 1.2 | 76.4 ± 1.1 | 65.9 ± 1.8 | 66.2 ± 2.1 | 67.7 ± 0.1 | 69.5 ± 1.4 | 64.8 ± 2.7 | 65.0 ± 3.3 |
| **QGP-1** | 61.1 ± 6.7 | 74.7 ± 9.0 | 65.5 ± 13.8 | 77.4 ± 3.8 | 79.4 ± 6.4 | 76.5 ± 4.4 | 82.7 ± 2.3 | 79.6 ± 6.0 |
| **LCC-18** | 57.8 ± 3.7 | 69.9 ± 2.0 | 65.5 ± 4.8 | 69.6 ± 3.9 | 64.3 ± 6.1 | 64.1 ± 3.3 | 68.7 ± 4.6 | 64.1 ± 3.3 |
| **H727** | 61.9 ± 2.8 | 61.1 ± 3.1 | 59.5 ± 0.6 | 61.3 ± 2.5 | 57.2 ± 2.3 | 60.6 ± 1.0 | 58.0 ± 0.8 | 63.7 ± 3.4 |
| **UMC-11** | 67.0 ± 0.2 | 72.5 ± 1.7 | 64.3 ± 3.7 | 79.1 ± 1.8 | 69.3 ± 2.3 | 73.4 ± 7.4 | 69.8 ± 0.2 | 74.1 ± 0.5 |

To investigate how key intracellular effectors were influenced by mTOR inhibitor treatment, all five NEN cell lines were analyzed using Western blot. p70S6 kinase, involved in cell growth and cell cycle progression, is known to be directly activated by mTOR. Consequently, mTOR inhibition should lead to decreased p70S6 kinase activation/phosphorylation. Indeed, this was found in lysates of all five NEN cell lines after incubation with 1 nM or 100 nM everolimus for 6 or 24 h (Figure 4). Yet, phospho-Akt levels also increased under mTOR inhibition, especially in BON and H727 cells. In UMC-11, phospho-Akt was barely detected, in QGP-1 cells it was very low, although total Akt was present at similar levels. Remarkably, the lower concentration of everolimus tended to lead to higher phospho-Akt levels than the higher concentration. Phosphorylation status of ERK1/2 was not affected in most cell lines; only BON showed slightly increased levels after treatment with everolimus. Neither caspase-3 nor PARP, markers for apoptosis, were cleaved under these conditions.

**Figure 4:**
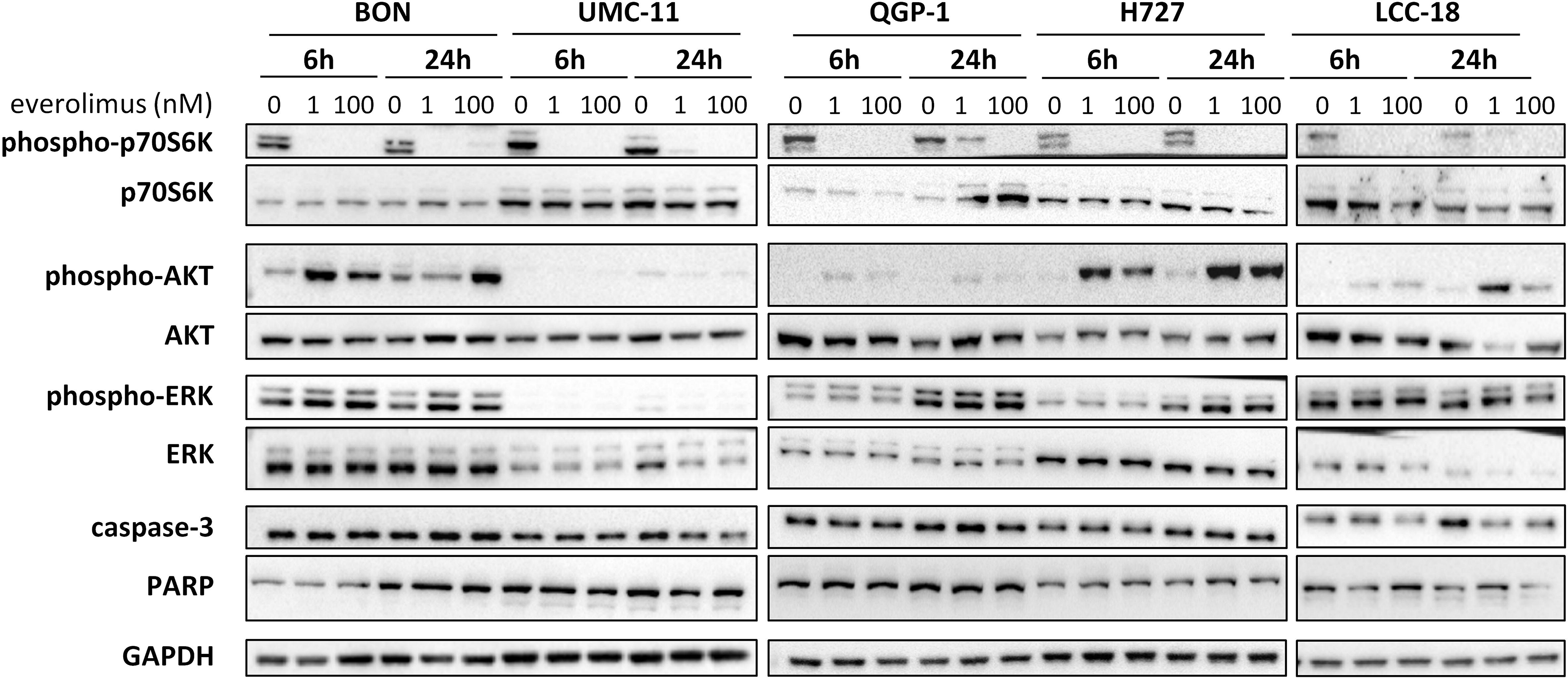
Effect of everolimus on intracellular signaling molecules. NEN cell lines were treated with vehicle (0 nM) or everolimus (1 nM, 100 nM) and harvested after 6 h and 24 h. Equal amounts of protein were separated on SDS gels and the indicated molecules detected after western blotting.

In summary, both mTOR inhibitors showed antiproliferative activity in all five NEN cell lines, caused by suppression of the mTOR signaling pathway and G1 cell cycle arrest. However, the cell lines differed in their reaction towards treatment, depending on counteracting regulation and the extent and duration of G1 cell accumulation. Whereas the rather resistant BON cells exhibited only a transient G1 arrest, but increased phospho-Akt levels, the most sensitive UMC-11 cells showed persistent G1 arrest and barely detectable phosphorylation of Akt.

### Radiation-induced effects on NEN cells

Before investigating combined mTOR inhibitor and radiation treatment, the sole impact of irradiation on NEN cells was assessed by applying a single dose of 0-10 Gy. Cell viability assays revealed a dose-dependent reduction of metabolic activity and cell number in all investigated NEN cell lines (Figure 5A). Cell numbers showed a more prominent reduction (60-80%) than metabolic activity (20-40%). The dose required to lower cell numbers by 50 % was approximately 8 Gy for H727, 6 Gy for BON and QGP-1, and 4 Gy for UMC-11 and LCC-18. In contrast to the observed G1 cell cycle arrest after mTOR inhibitor treatment, irradiation with 10 Gy resulted in a strong accumulation of cells in G2/M cell cycle phase after 24 h (Figure 5B). QGP-1 cells reached the highest increase (55%), followed by BON (48 %), LCC-18 (38%), H727 (31 %) and UMC-11 (31 %) (**Table 3**). This G2/M cell cycle arrest was retained over time until 96 h after irradiation in all cell lines, although percentages partly decreased (Figure S1).

**Figure 5:**
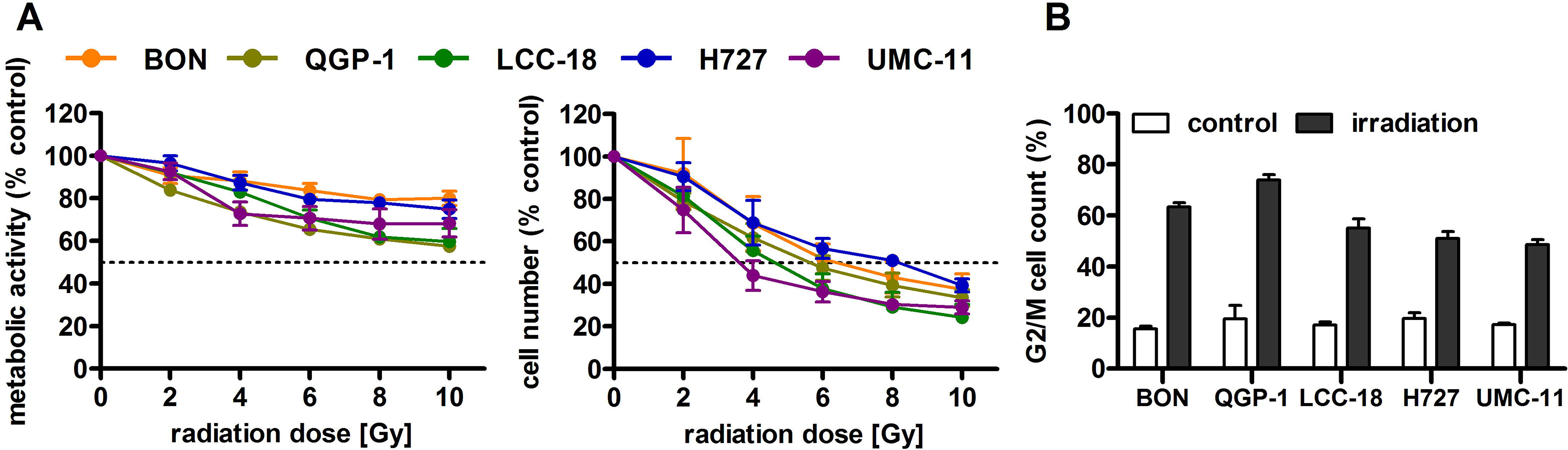
Irradiation leads to reduced cell viability and G2/M accumulation of NEN cells. **(A)** NEN cell lines were irradiated with doses of 0 to 10 Gy, incubated for 96 h and analyzed for metabolic activity and cell number. Data represent mean ± S.E.M. (n=2-3). **(B)** NEN cell lines were irradiated with 10 Gy and samples were collected after 24 h for cell cycle analysis by flow cytometry. Data are shown as mean ± S.E.M. (n=2-3).

**Table 3.** Percentages of cells in G2/M cell cycle after irradiation. Data show mean ± S.E.M. (n=2-3).

|  | 24h |  | 48h |  | 72h |  | 96h |  |
| --- | --- | --- | --- | --- | --- | --- | --- | --- |
|  | 0 | 10 Gy | 0 | 10 Gy | 0 | 10 Gy | 0 | 10 Gy |
| **BON** | 15.6 ± 1.0 | 63.3 ± 1.6 | 16.3 ± 0.4 | 54.5 ± 2.4 | 17.0 ± 0.6 | 41.7 ± 2.5 | 17.0 ± 1.5 | 38.1 ± 0.6 |
| **QGP-1** | 19.4 ± 5.3 | 73.9 ± 2.1 | 18.7 ± 7.3 | 53.0 ± 3.2 | 9.9 ± 3.0 | 46.9 ± 3.0 | 8.9 ± 1.2 | 43.2 ± 1.9 |
| **LCC-18** | 17.0 ± 1.2 | 55.0 ± 3.8 | 12.5 ± 2.7 | 48.0 ± 0.9 | 11.5 ± 1.2 | 42.8 ± 2.2 | 10.0 ± 1.6 | 33.9 ± 0.3 |
| **H727** | 19.6 ± 2.2 | 50.9 ± 2.8 | 19.8 ± 1.5 | 52.4 ± 1.4 | 20.1 ± 1.2 | 45.1 ± 1.6 | 21.1 ± 2.0 | 37.9 ± 2.1 |
| **UMC-11** | 17.1 ± 0.7 | 48.5 ± 2.0 | 19.1 ± 2.0 | 62.7 ± 1.1 | 15.5 ± 0.6 | 51.2 ± 3.3 | 13.9 ± 0.9 | 47.5 ± 2.6 |

### Combined effect of temsirolimus and irradiation

The five NEN cell lines under investigation were pretreated with temsirolimus for 24 h, subsequently irradiated and incubated for another 96 h. As observed before, cell numbers declined when applying increasing doses of temsirolimus or radiation (Figure 6). However, combination of both reduced cell numbers to a greater extent, especially in the low nanomolar range. UMC-11 cells were already dramatically affected by temsirolimus alone; therefore, the additive effect of irradiation was rather moderate when compared to BON, QGP-1 or LCC-18. The biphasic inhibition pattern of temsirolimus was retained after irradiation, although the curve slopes flattened out with increasing radiation doses (Figure 6). For a detailed analysis, Figure 7 shows the results for the combination of temsirolimus at 1 nM or 1 μM with 4 Gy of irradiation. For both concentrations, sequential treatment resulted in a higher reduction of cell numbers than the respective single treatments. In the case of pretreatment with 1 μM, cell numbers for BON and UMC-11 decreased significantly when compared to irradiation alone (Figure 7B).

**Figure 6:**
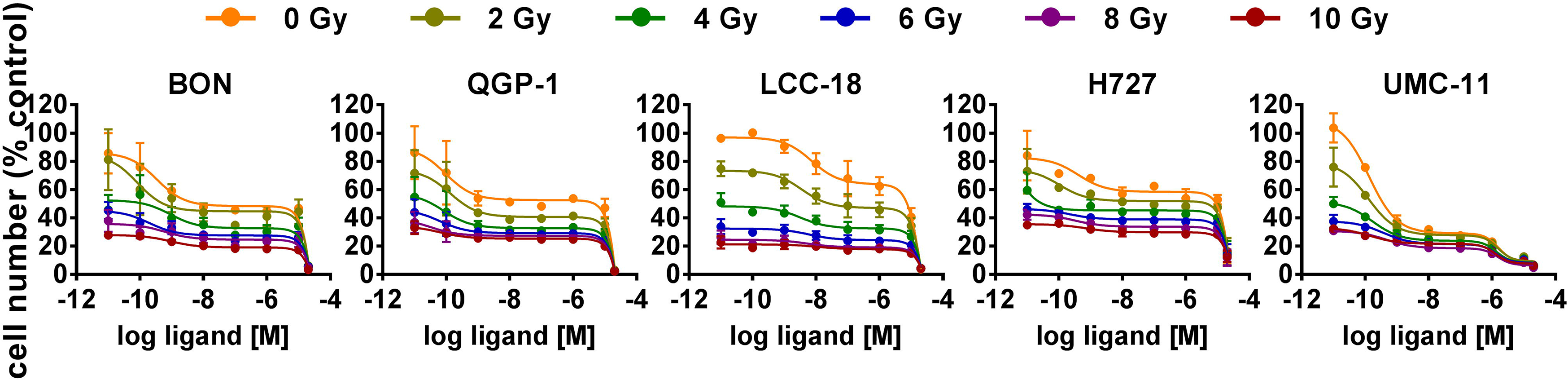
Additive effect of temsirolimus and irradiation on NEN cell numbers. NEN cell lines were pretreated with increasing concentrations of temsirolimus (0.01 nM to 20 μM) for 24 h before irradiation. Cell number was determined 96 h after irradiation with different doses of 0 to 10 Gy. Graphs show mean ± S.E.M. (n=2-3).

**Figure 7:**
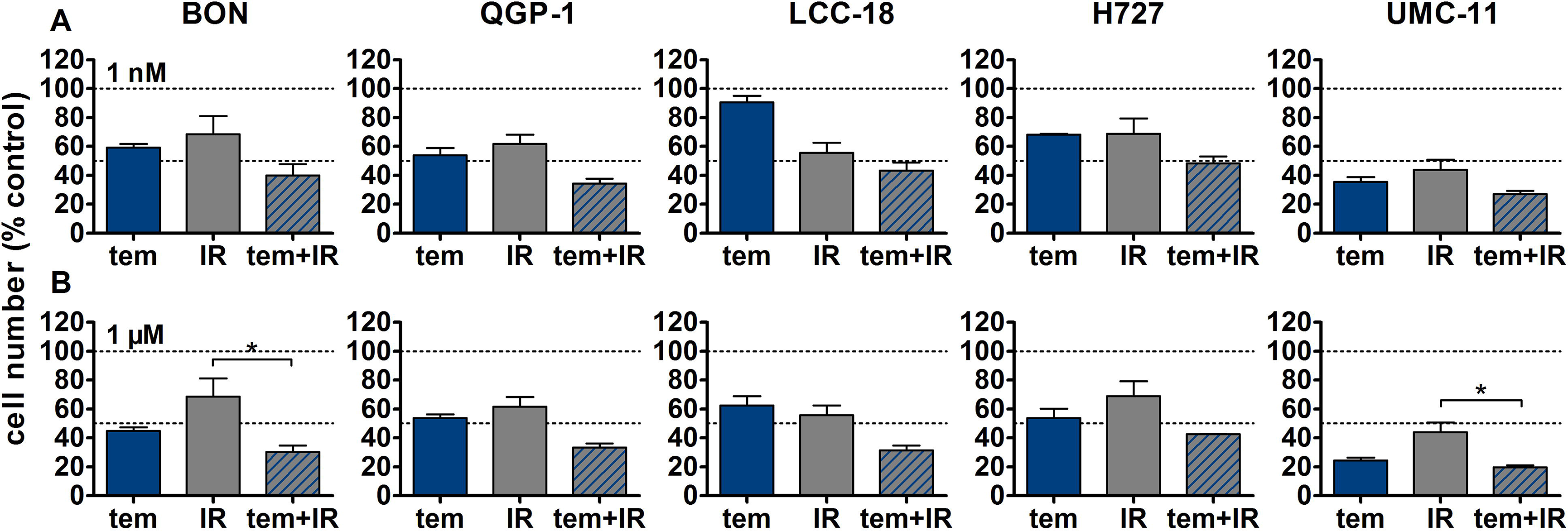
Detailed analysis of cell numbers for combined treatment of temsirolimus and 4 Gy. NEN cell lines were pretreated with 1 nM **(A)**or 1 μM **(B)** temsirolimus for 24 h before irradiation with 4 Gy. Cell number was determined 96 h after irradiation. Graphs show mean ± S.E.M. (n=2-3). Evaluated with one-way ANOVA (Kruskal-Wallis test) and Dunn’s posttest, * P≤0.05. tem, temsirolimus; IR, irradiation.

In addition, clonogenic survival assays were included to evaluate the impact of the combined treatment over a longer period. For this, NEN cells were irradiated with 0-4 Gy after preincubation with temsirolimus. Combination of both interventions resulted in clearly impaired cell survival in comparison to the single treatments (Figure 8). Radiation doses higher than 4 Gy resulted in barely visible colonies in the wells (data not shown). This is in strong contrast to the shorter cell number assays, where the highest dose of 10 Gy still resulted in detectable signals (Figure 6). Irradiation with 4 Gy alone inhibited cell numbers only by 32 % (BON), 38 % (QGP-1), 44 % (LCC-18), 31 % (H727) and 56 % (UMC-11) (Figure 7). Cell survival in the clonogenic assay on the other hand was lowered by 45 % (BON), 55 % (QGP-1), 13 % (LCC-18), 75 % (H727) and 92 % (UMC-11) (Figure). In BON and H727, pretreated with 1 nM temsirolimus, and in QGP-1, pretreated with 1 μM, additional irradiation with 4 Gy significantly decreased cell survival in comparison to temsirolimus alone (Figure). The beneficial impact of a combined treatment regimen was seen in all cell lines and lowered cell survival down to ≤ 20 %.

**Figure 8:**
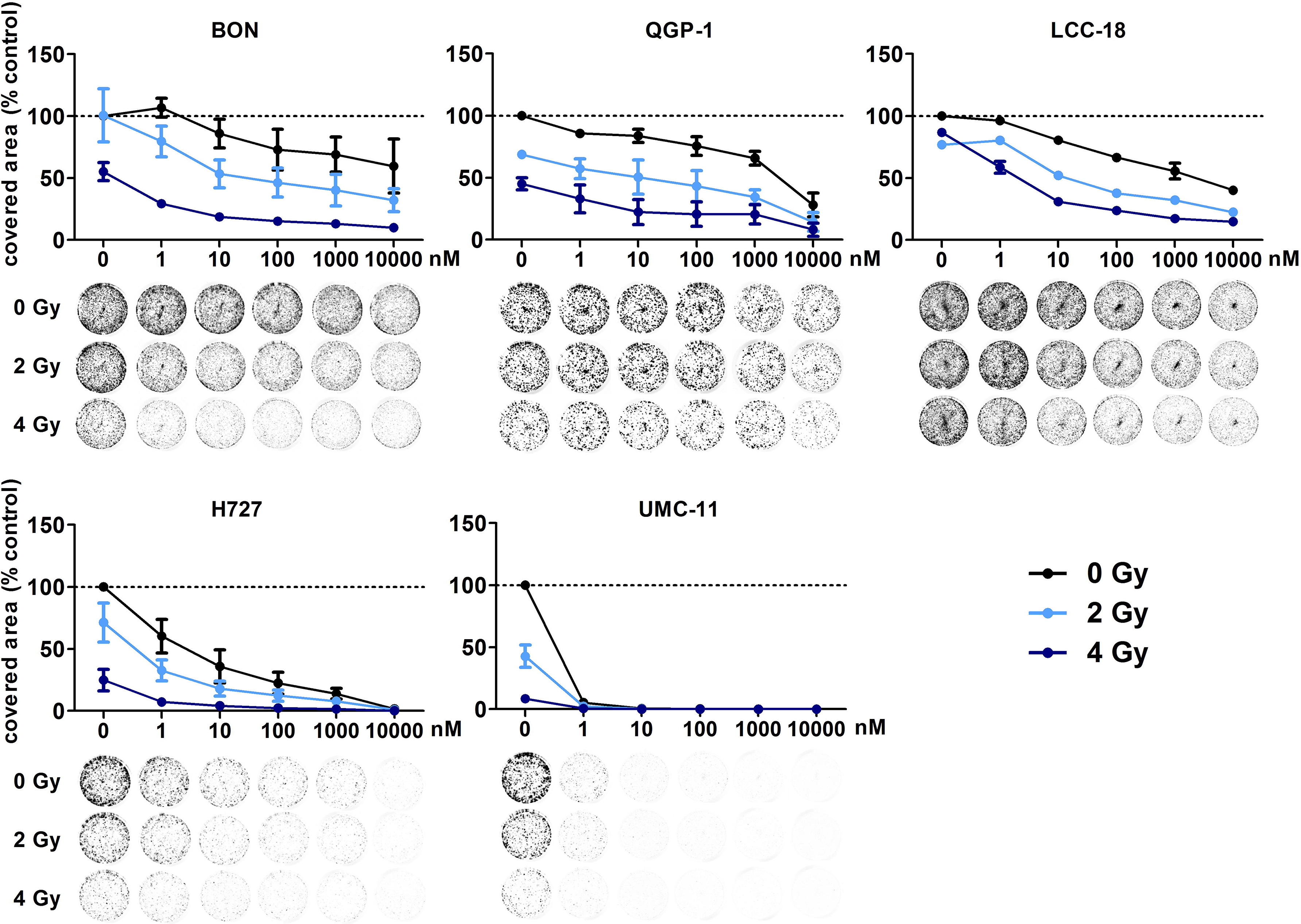
Additive effect of temsirolimus and irradiation on NEN cell survival. NEN cell lines were seeded at low density, treated with increasing concentrations of temsirolimus for 24 h before irradiation with 0-4 Gy. Cells were incubated for 1-2 weeks until colony formation. Data were normalized to untreated controls and represent mean ± S.E.M. (n=2-3).

**Figure 9:**
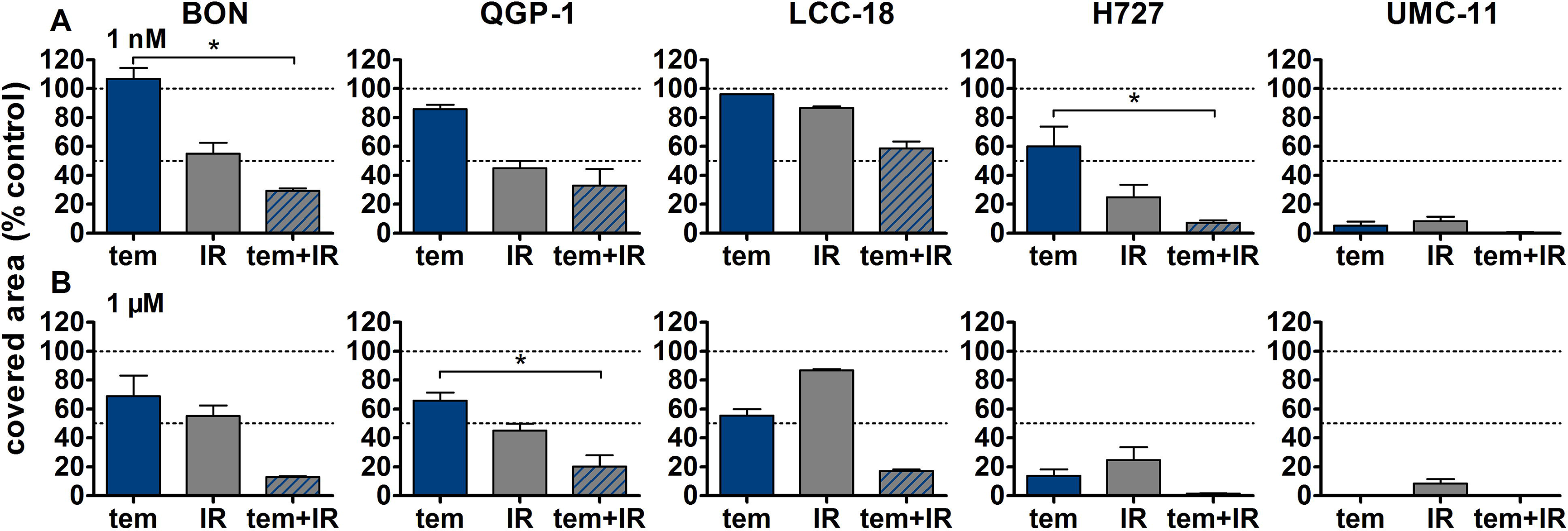
Detailed analysis of cell survival for combined treatment of temsirolimus and 4 Gy. NEN cell lines were pretreated with 1 nM **(A)** or 1 μM **(B)** temsirolimus for 24 h before irradiation with 4 Gy. Cell survival was determined 1-2 weeks after irradiation. Graphs show mean ± S.E.M. (n=2-3). Evaluated with one-way ANOVA (Kruskal-Wallis test) and Dunn’s posttest, * P≤0.05. tem, temsirolimus; IR, irradiation.

**Figure 10:**
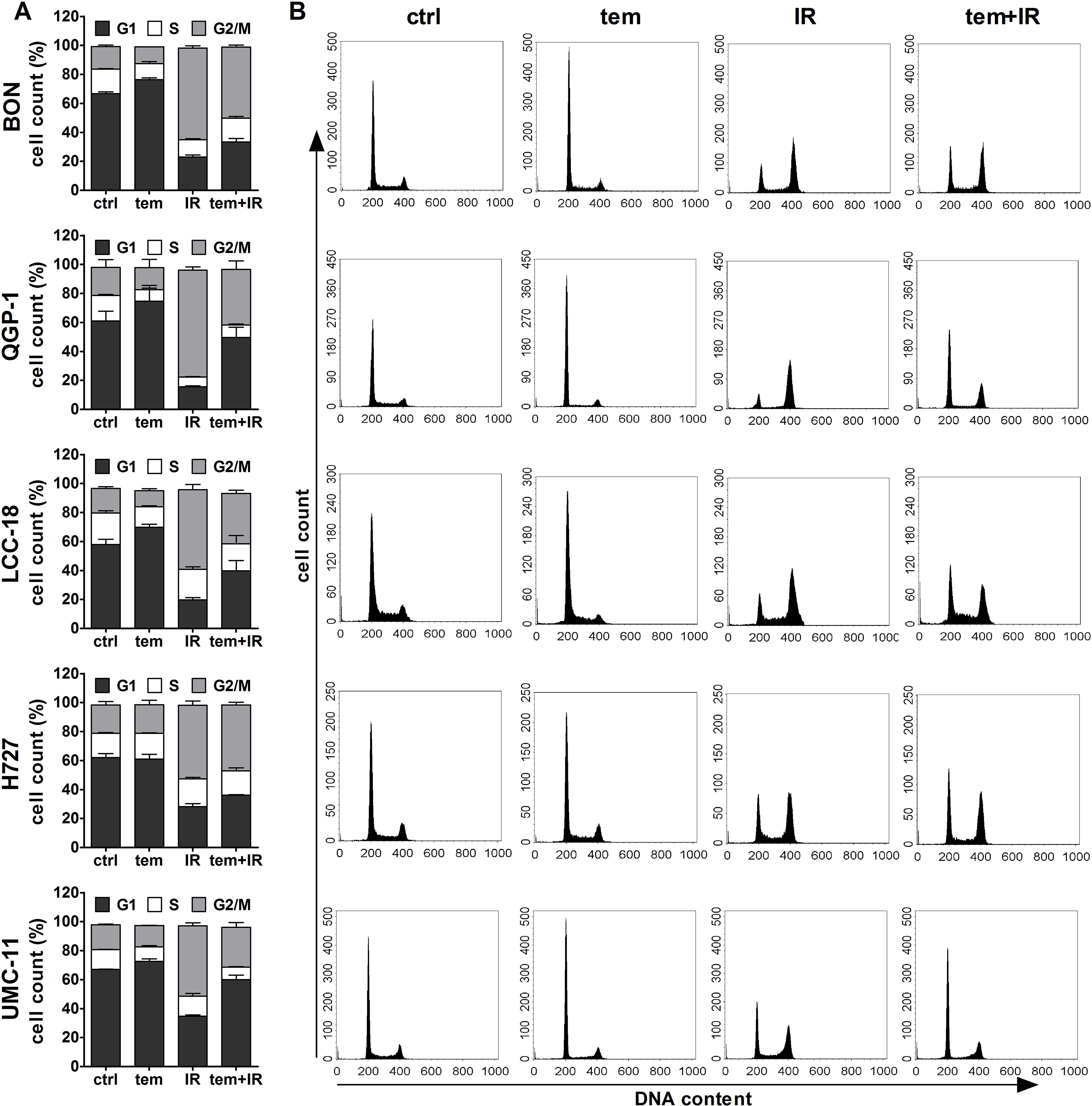
Temsirolimus pretreatment abrogates radiation-induced G2/M cell accumulation. NEN cell lines were incubated with 1 μM temsirolimus (tem) or vehicle (ctrl) for 24 h before irradiation. For assessment of cell cycle distribution pretreated NEN cells were collected 24 h after irradiation with 10 Gy (IR, tem+IR), stained with propidium iodide and analyzed by flow cytometer. Data are shown as bar diagrams with mean ± S.E.M. (n=2-3) **(A)** or as DNA histograms of one representative experiment **(B)**.

The previous analysis of cell cycle distribution following the respective single treatments revealed increased cell accumulation in G1 after mTOR inhibitor treatment, or in G2/M after irradiation. Here, the combined effect of both treatments was evaluated by preincubating the respective cell lines with 1 μM temsirolimus for 24 h before irradiating them with 10 Gy. In all five NEN cell lines, mTOR inhibitor pretreatment resulted in a diminished radiation-induced G2/M arrest after 24 h (Figure). Correspondingly, the G1 fraction increased. In QGP-1 cells, the impact of combined treatment on cell cycle distribution was most profound. Compared to irradiation only, the G2/M fraction decreased by 49 % (from 74 to 38 %), while cells in G1 increased by 212 % (from 16 to 50 %) (Table 4). Similarly, UMC-11, LCC-18 and BON cells showed a decrease in G2/M and an increase in G1. Even in H727 cells, which did not show any G1 cell accumulation when applying temsirolimus alone, combined treatment led to an increase of the G1 fraction after 24 h. This reduced G2/M arrest after combinatorial treatment was maintained for up to 96 h in UMC-11 and H727 (Figure S2), whereas it leveled out after 48 h in BON and LCC-18 cells. QGP-1 cells slowly returned to the irradiation-like cell cycle distribution with complete adjustment after 96 h. The sub-G1 fraction as a measure for nuclear debris was increased after irradiation only or after combination with temsirolimus pretreatment in all investigated cell lines.

**Table 4.** Percentages of cells in G1 and G2/M cell cycle 24 h after irradiation. Data show mean ± S.E.M. (n=2-3).

|  | G1 |  |  |  | G2/M |  |  |  |
| --- | --- | --- | --- | --- | --- | --- | --- | --- |
|  | ctrl | tem | IR | tem + IR | ctrl | tem | IR | tem + IR |
| **BON** | 66.7 ± 1.2 | 76.4 ± 1.1 | 22.9 ± 1.5 | 33.4 ± 2.4 | 15.6 ± 1.0 | 11.5 ± 0.0 | 63.3 ± 1.6 | 49.1 ± 1.5 |
| **QGP-1** | 61.1 ± 6.7 | 74.7 ± 9.0 | 15.6 ± 0.6 | 49.5 ± 7.2 | 19.4 ± 5.3 | 15.2 ± 5.7 | 73.9 ± 2.1 | 38.4 ± 5.9 |
| **LCC-18** | 57.8 ± 3.7 | 69.9 ± 2.0 | 19.7 ± 1.5 | 39.7 ± 7.1 | 17.0 ± 1.2 | 11.0 ± 1.5 | 55.0 ± 3.8 | 34.8 ± 2.2 |
| **H727** | 61.9 ± 2.8 | 61.1 ± 3.1 | 28.2 ± 1.9 | 36.3 ± 0.2 | 19.6 ± 2.2 | 19.8 ± 3.1 | 50.9 ± 2.8 | 45.3 ± 1.9 |
| **UMC-11** | 67.0 ± 0.2 | 72.5 ± 1.7 | 34.7 ± 0.8 | 59.9 ± 3.2 | 17.1 ± 0.7 | 14.7 ± 0.1 | 48.5 ± 2.0 | 27.4 ± 3.3 |

In summary, combination of mTOR inhibitor treatment and irradiation indicated superiority compared to the single treatments as proven by increased decrease of cell numbers and cell survival. Preincubation with temsirolimus abrogated the irradiation-induced G2/M arrest in all NEN cell lines under investigation.

## Discussion

PRRT has become an essential treatment option for advanced NEN. It demonstrated increased progression-free survival when compared to chemotherapy, radiation and intervention with somatostatin analogs (SSAs). Objective response rates range between 15 and 35 % (24). However, complete remissions are still very rare, although PRRT delivers high doses specifically to the tumor (3). In contrast, similar nuclear therapies such as radioiodine therapy for thyroid cancers or radioimmunotherapy for B-cell non-Hodgkin’s lymphoma achieve complete remissions in 30 % or 75 % of all cases, respectively (25,26). It was hypothesized that other established NEN therapies such as mTOR inhibitors may have a radiosensitizing effect, leading to a beneficial impact on the outcome of PRRT. Therefore, this work evaluated the radiosensitizing potential of mTOR inhibitor treatment prior to irradiation in a panel of five NEN cell lines of different origin.

First, the impact of the two mTOR inhibitors temsirolimus and everolimus on cellular processes was validated for the utilized NEN cell lines. Proliferation assays revealed a biphasic inhibition of metabolic activity and cell number, with two resulting IC_50_ values for both substances in all five cell lines. Whereas nanomolar concentrations led to a moderate antiproliferative effect, micromolar concentrations suppressed cell viability completely. This is in line with a report from Shor et al. that also described a low-dose and high-dose effect of temsirolimus in different cancer cell lines (27). In addition to the well-known FKBP12-dependent binding of temsirolimus to mTOR, they suggest a second FKBP12 independent mechanism, which might be responsible for the profound high dose effect. Although the determined low-dose IC_50_ values were approximately 1 nM for all cell lines and both inhibitors, efficacy (amplitude of the effect) differed between the investigated cell lines. Pulmonary UMC-11 cells were found to be highly drug-sensitive, as cell viability was reduced by 75 % after treatment at nanomolar concentrations, whereas BON cells proved to be rather resistant (20 % reduction). Long-term clonogenic survival assays over up to two weeks confirmed these results.

Signaling and cell cycle analysis were conducted to further elucidate the observed response in these cell lines. Similar to a study by Hurvitz et al. in breast cancer cell lines (28), inhibition of p70S6 kinase, a downstream target of mTOR, was achieved in all NEN cell lines, but did not correlate with their susceptibility as measured by cell viability and survival. Yet, phospho-p70S6 kinase levels were only monitored for 24 h after treatment and later time points might reveal divergent response patterns. The high sensitivity of UMC-11 might be explained by a constantly elevated fraction of cells in the G1 cell cycle phase, up to 96 h after treatment. In addition, mTOR inhibition did not result in a counteracting phospho-Akt upregulation in these cells – a frequently observed resistance mechanism in tumor cells based on loss of the negative S6K feedback loop on IRS and PI3K. In comparison, the rather resistant BON cells show only transient G1 cell accumulation of up to 24 to 48 h and a clear increase of phospho-Akt and phospho-ERK levels, indicating several escape pathways. This is in line with a report by Zitzmann et al., that investigated the effect of everolimus on proliferation, cell cycle distribution and signaling of BON cells (29). In contrast to their results, this study could not confirm apoptosis induction in these cells, though investigating similar time points and concentrations. Interestingly, between these very distinct cases of BON and UMC-11 cells, intermediate phenotypes seem to exist. For example, despite a clear Akt activation and a rather weak and transient G1 cell accumulation, clonogenic survival of H727 was strongly reduced after mTOR inhibition. QGP-1 cells behaved similar to BON cells (both expressing mutant p53) with regard to cell viability and survival assays, with a transient but very strong G1 cell accumulation. On the other hand, QGP-1 cells did not activate Akt upon mTOR inhibition.

Cell cycle checkpoints at G1/S or G2/M phase transitions are important control mechanisms to maintain genomic integrity within the cell in response to environmental stress and DNA damaging agents such as irradiation. Their activation is mediated by a complex signaling network including cyclins, cyclin-dependent kinases and the key modulator p53, which is primarily involved in G1 cell cycle arrest (30). In the case of extensive and irreparable damage, cells go into apoptosis. It has been shown that mTOR inhibitors induce a p53-independent G1 cell cycle arrest by an impaired translation of cyclin D1 and an enhanced expression of p27Kip1 (31–33). Neither molecule was included in this analysis, but differences in their expression upon mTOR inhibition might further explain the varying treatment susceptibility of the single cell lines. The permanent G1 cell cycle accumulation of drug-sensitive UMC-11 cells indicates a likely influence of the cells’ capability to stay in this arrest over time.

Several genetic alterations have been associated with mTOR sensitivity such as PTEN-deficiency (34), a distinct SNP in FGFR4 (35) or PIK3CA mutations (36). Undoubtedly, an increased basal activation of mTOR and its signaling pathway plays an important role for the outcome of mTOR inhibitor treatment. Nevertheless, clinical biomarkers for a reliable response prediction were not identified so far (37). In conclusion, this study verifies the expected effects of mTOR inhibitors in vitro and further complements the reported results for pancreatic neuroendocrine BON and QGP-1 cells (29,34) by evaluating an extended NEN cell line panel that includes pulmonary neuroendocrine H727 and UMC-11 as well as colonic neuroendocrine LCC-18 cells.

In line with many reports that analyzed radiation effects and DNA damage responses in cancer cells, all investigated NEN cell lines revealed an accumulation at the G2/M junction, which was retained over time. Radiation susceptibility differed only slightly between cell lines as determined by cell counting. Although DNA-damaging radiation is primarily associated with G1 arrest, it was postulated that most cancer cells lack a functional G1 checkpoint due to mutations in the key molecule p53. Therefore, they are more reliant on the pre-mitotic G2/M checkpoint for repair of potentially lethal damage and display a strong G2/M arrest upon irradiation (38–40). Cell viability and survival assays revealed the superiority of combining mTOR inhibitors with irradiation in comparison to either single application. In all NEN cell lines under investigation, this treatment strategy exhibited an additive inhibitory effect. Interestingly, the response of the drug-sensitive drug-sensitive UMC-11 cells was barely enhanced by this approach as mTOR inhibition already impaired survival to a great extent. As discussed before, mTOR inhibitors induced a G1 cell cycle accumulation, whereas after irradiation cells accumulated in G2/M. In combination, pretreatment with temsirolimus clearly diminished radiation-induced G2/M arrest in all five NEN cell lines. Thus, it can be hypothesized that temsirolimus prevents DNA damage repair processes that normally occur during G2/M arrest. Thereby, cells with unrepaired DNA lesions may prematurely enter mitosis and undergo the so called mitotic catastrophe, which is distinct from apoptotic cell death (41,42). Other groups addressing the radiosensitizing effect of everolimus consistently reported an enhanced inhibition of cell growth in vitro and in vivo when applying the combinatorial treatment regimen. However, cell cycle distribution analyses revealed different outcomes. Su et al. observed no difference in G2/M between everolimus with and without radiation in Ras-transformed cells (43). Possibly, the chosen concentration of 30 nM was too low to uncover the G2/M abrogation. In contrast, Nassim et al. reported an increase of both G1 and G2/M when combining everolimus with radiation in bladder cancer cells (44). They suggest that the everolimus-induced G1 arrest confers an enhanced sensitivity for the following radiation as fewer cells were counted in the rather radioresistant S-phase (45). However, in NEN cell lines the proportion of S-phase cells was relatively unaffected. It must be further noted, that the length of pretreatment differed between the studies, which might have a significant impact on the evaluated parameters. In line with the presented work, Wang et al. reported an abrogation of the radiation-induced G2/M arrest in pancreatic cells, but after pretreatment with metformin (46).

One of the limitations of this study is the use of external beam irradiation instead of radioligands used in PRRT, such as ^177^Lu-DOTATOC or ^177^Lu-DOTATATE. The external beam irradiation used was from a calibrated ^137^Cs source. It delivers reproducible, stable and precise dose levels to cultured cells and is easily accessible. Similar to ^177^Lu, ^137^Cs is a combined beta/gamma emitter. Though both isotopes have a similar beta emission energy, their effect on living cells will differ to a small extent. In contrast to external beam irradiation, PRRT with ^177^Lu-coupled SSA only targets cells expressing somatostatin receptors (SSTR), preferentially SSTR2. However, for all NEN cell lines under investigation in this study, we have previously demonstrated low target expression and a lack of binding of the SSA octreotide. All five cell lines cells showed 10- to 1000-fold lower SSTR2 mRNA levels than human NEN tissues (47). Consequently, external beam irradiation had to be utilized. In comparison to targeted PRRT with ^177^Lu-coupled peptides, this type of irradiation is undirected and affects cells independent of their receptor status. It would certainly be of great interest to see the experimental outcome in a human SSTR2-transfected NEN cell model or in primary patient-derived NEN cells. In addition to treatment with ^177^Lu-DOTATOC or ^177^Lu-DOTATATE, the evaluation of the antagonistic peptide ligand ^177^Lu-DOTA-JR11 could be an interesting part of such a combinatorial in vivo study. Recently, a favorable toxicity profile has been found for the combination of temsirolimus and ^177^Lu-DOTATATE in rats (48). In addition, further translational research is required to assess the potential of the combination of PRRT and mTOR inhibitors in pulmonary carcinoids as well as in G2 and G3 NEN. Likewise, careful preclinical evaluation in a suitable animal model is warranted, especially as the combination of ^177^Lu-DOTATATE and RAD001 had led to enhanced metastasis in a rat NEN model (49). The future application of somatostatin antagonists as single agents or together with mTOR inhibitors as well as other substances may provide new opportunities for SSTR-based therapies with improved response rates.

## Acknowledgements

The authors wish to thank Sarah Erdmann for helpful discussions in the early phase of this project.

## Disclosure Statement

The authors declare no potential conflicts of interest.

## Funding Sources and their Role

This work was supported by grants from the German Ministry of Education and Research (BMBF IP614 and IPT614A to Carsten Grötzinger). Samantha Exner was the recipient of a fellowship by Sonnenfeld-Stiftung. The authors acknowledge support from the German Research Foundation (DFG) and the Open Access Publication Fund of Charité – Universitätsmedizin Berlin. Funding agencies had no influence on design, data acquisition of evaluation in any part of this study.

## Author Contributions

CG and VP designed and supervised research; SE and GAT performed research and analyzed data; SE and CG wrote the paper.

## Notes

https://doi.org/10.5281/zenodo.3922212

### Competing Interest Statement

The authors have declared no competing interest.

## References

1. Kwekkeboom DJ, de Herder WW, Kam BL, van Eijck CH, van Essen M, Kooij PP, et al. Treatment with the radiolabeled somatostatin analog [177 Lu-DOTA 0,Tyr3]octreotate: toxicity, efficacy, and survival. J Clin Oncol. 2008 May;26(13):2124–30.

2. Strosberg J, El-Haddad G, Wolin E, Hendifar A, Yao J, Chasen B, et al. Phase 3 Trial of 177 Lu-Dotatate for Midgut Neuroendocrine Tumors. N Engl J Med. 2017 Jan;376(2):125–35.

3. Cremonesi M, Ferrari M, Bodei L, Tosi G, Paganelli G. Dosimetry in Peptide radionuclide receptor therapy: a review. J Nucl Med. 2006;47(9):1467–75.

4. Klümpen H-J, Beijnen JH, Gurney H, Schellens JHM. Inhibitors of mTOR. Oncologist. 2010 Jan;15(12):1262–9.

5. Laplante M, Sabatini DM. mTOR signaling in growth control and disease. Cell. 2012 Apr;149(2):274–93.

6. Sabatini DM. mTOR and cancer: insights into a complex relationship. Nat Rev Cancer. 2006 Sep;6(9):729–34.

7. O’Reilly KE, Rojo F, She Q-B, Solit D, Mills GB, Smith D, et al. mTOR inhibition induces upstream receptor tyrosine kinase signaling and activates Akt. Cancer Res. 2006 Feb;66(3):1500–8.

8. Sarbassov DD, Ali SM, Sengupta S, Sheen J-H, Hsu PP, Bagley AF, et al. Prolonged rapamycin treatment inhibits mTORC2 assembly and Akt/PKB. Mol Cell. 2006 Apr;22(2):159–68.

9. Neshat MS, Mellinghoff IK, Tran C, Stiles B, Thomas G, Petersen R, et al. Enhanced sensitivity of PTEN-deficient tumors to inhibition of FRAP/mTOR. Proc Natl Acad Sci U S A. 2001 Aug;98(18):10314–9.

10. Carracedo A, Ma L, Teruya-Feldstein J, Rojo F, Salmena L, Alimonti A, et al. Inhibition of mTORC1 leads to MAPK pathway activation through a PI3K-dependent feedback loop in human cancer. J Clin Invest. 2008 Sep;118(9):3065–74.

11. Lee MS, O’Neil BH. Summary of emerging personalized medicine in neuroendocrine tumors: are we on track? J Gastrointest Oncol. 2016 Oct;7(5):804–18.

12. Yao JC, Shah MH, Ito T, Bohas CL, Wolin EM, Van Cutsem E, et al. Everolimus for advanced pancreatic neuroendocrine tumors. N Engl J Med. 2011 Feb;364(6):514–23.

13. Yao JC, Fazio N, Singh S, Buzzoni R, Carnaghi C, Wolin E, et al. Everolimus for the treatment of advanced, non-functional neuroendocrine tumours of the lung or gastrointestinal tract (RADIANT-4): a randomised, placebo-controlled, phase 3 study. Lancet (London, England). 2016 Mar;387(10022):968–77.

14. Pavel ME, Hainsworth JD, Baudin E, Peeters M, Hörsch D, Winkler RE, et al. Everolimus plus octreotide long-acting repeatable for the treatment of advanced neuroendocrine tumours associated with carcinoid syndrome (RADIANT-2): a randomised, placebo-controlled, phase 3 study. Lancet. 2011;378(9808):2005–12.

15. Yao JC, Lombard-Bohas C, Baudin E, Kvols LK, Rougier P, Ruszniewski P, et al. Daily oral everolimus activity in patients with metastatic pancreatic neuroendocrine tumors after failure of cytotoxic chemotherapy: a phase II trial. J Clin Oncol. 2010 Jan;28(1):69–76.

16. Eads JR, Meropol NJ. A new era for the systemic therapy of neuroendocrine tumors. Oncologist. 2012;17(3):326–38.

17. Duran I, Kortmansky J, Singh D, Hirte H, Kocha W, Goss G, et al. A phase II clinical and pharmacodynamic study of temsirolimus in advanced neuroendocrine carcinomas. Br J Cancer. 2006 Nov;95(9):1148–54.

18. Atkins MB, Yasothan U, Kirkpatrick P. Everolimus. Nat Rev Drug Discov. 2009 Jul;8(7):535–6.

19. Rini B, Kar S, Kirkpatrick P. Temsirolimus. Nat Rev Drug Discov. 2007 Aug;6:599.

20. Claringbold PG, Turner JH. NeuroEndocrine Tumor Therapy with Lutetium-177-octreotate and Everolimus (NETTLE): A Phase I Study. Cancer Biother Radiopharm. 2015 Aug;30(6):261–9.

21. Krishan A. Rapid flow cytofluorometric analysis of mammalian cell cycle by propidium iodide staining. J Cell Biol. 1975 Jul;66(1):188–93.

22. Wersto RP, Chrest FJ, Leary JF, Morris C, Stetler-Stevenson MA, Gabrielson E. Doublet discrimination in DNA cell-cycle analysis. Cytometry. 2001 Oct;46(5):296–306.

23. Guzmán C, Bagga M, Kaur A, Westermarck J, Abankwa D. ColonyArea: An ImageJ Plugin to Automatically Quantify Colony Formation in Clonogenic Assays. Rota R, editor. PLoS One. 2014 Mar;9(3):e92444.

24. Kwekkeboom DJ, Krenning EP. Peptide Receptor Radionuclide Therapy in the Treatment of Neuroendocrine Tumors. Hematol Oncol Clin North Am. 2016 Feb;30(1):179–91.

25. Goldsmith SJ. Radioimmunotherapy of lymphoma: Bexxar and Zevalin. Semin Nucl Med. 2010 Mar;40(2):122–35.

26. Luster M, Hänscheid H, Freudenberg LS, Verburg FA. Radioiodine therapy of metastatic lesions of differentiated thyroid cancer. J Endocrinol Invest. 2012;35(6 Suppl):21–9.

27. Shor B, Zhang W-G, Toral-Barza L, Lucas J, Abraham RT, Gibbons JJ, et al. A new pharmacologic action of CCI-779 involves FKBP12-independent inhibition of mTOR kinase activity and profound repression of global protein synthesis. Cancer Res. 2008 Apr;68(8):2934–43.

28. Hurvitz SA, Kalous O, Conklin D, Desai AJ, Dering J, Anderson L, et al. In vitro activity of the mTOR inhibitor everolimus, in a large panel of breast cancer cell lines and analysis for predictors of response. Breast Cancer Res Treat. 2015 Feb;149(3):669–80.

29. Zitzmann K, De Toni EN, Brand S, Göke B, Meinecke J, Spöttl G, et al. The novel mTOR inhibitor RAD001 (everolimus) induces antiproliferative effects in human pancreatic neuroendocrine tumor cells. Neuroendocrinology. 2007;85(1):54–60.

30. Cooper G. The Cell: A Molecular Approach. 2nd ed. Sunderland (MA): Sinauer Associates; The Eukaryotic Cell Cycle.; 2000.

31. Grewe M, Gansauge F, Schmid RM, Adler G, Seufferlein T. Regulation of cell growth and cyclin D1 expression by the constitutively active FRAP-p70s6K pathway in human pancreatic cancer cells. Cancer Res. 1999 Aug;59(15):3581 LP – 3587.

32. Kawamata S, Sakaida H, Hori T, Maeda M, Uchiyama T. The upregulation of p27Kip1 by rapamycin results in G1 arrest in exponentially growing T-cell lines. Blood. 1998 Jan;91(2):561 LP – 569.

33. Hashemolhosseini S, Nagamine Y, Morley SJ, Desrivie S, Mercep L, Ferrari S. Rapamycin Inhibition of the G 1 to S Transition Is Mediated by Effects on Cyclin D1 mRNA and Protein Stability. J Biol Chem. 1998;273(23):14424–9.

34. Missiaglia E, Dalai I, Barbi S, Beghelli S, Falconi M, della Peruta M, et al. Pancreatic Endocrine Tumors: Expression Profiling Evidences a Role for AKT-mTOR Pathway. J Clin Oncol. 2010 Jan;28(2):245–55.

35. Serra S, Zheng L, Hassan M, Phan AT, Woodhouse LJ, Yao JC, et al. The FGFR4-G388R single-nucleotide polymorphism alters pancreatic neuroendocrine tumor progression and response to mTOR inhibition therapy. Cancer Res. 2012 Nov;72(22):5683–91.

36. Weigelt B, Warne PH, Downward J. PIK3CA mutation, but not PTEN loss of function, determines the sensitivity of breast cancer cells to mTOR inhibitory drugs. Oncogene. 2011 Jul;30(29):3222–33.

37. Zatelli MC, Fanciulli G, Malandrino P, Ramundo V, Faggiano A, Colao A, et al. Predictive factors of response to mTOR inhibitors in neuroendocrine tumours. Endocr Relat Cancer. 2016 Mar;23(3):R173–83.

38. Bucher N, Britten CD. G2 checkpoint abrogation and checkpoint kinase-1 targeting in the treatment of cancer. Br J Cancer. 2008 Feb;98(3):523–8.

39. DiPaola RS. To arrest or not to G(2)-M Cell-cycle arrest : commentary re: A. K. Tyagi et al., Silibinin strongly synergizes human prostate carcinoma DU145 cells to doxorubicin-induced growth inhibition, G(2)-M arrest, and apoptosis. Clin. cancer res., 8: 3512-3519, 2. Clin Cancer Res. 2002 Nov;8(11):3311–4.

40. Chen T, Stephens PA, Middleton FK, Curtin NJ. Targeting the S and G2 checkpoint to treat cancer. Drug Discov Today. 2012 Mar;17(5–6):194–202.

41. Vitale I, Galluzzi L, Castedo M, Kroemer G. Mitotic catastrophe: a mechanism for avoiding genomic instability. Nat Rev Mol Cell Biol. 2011 Jun;12(6):385–92.

42. Kawabe T. G2 checkpoint abrogators as anticancer drugs. Mol Cancer Ther. 2004 Apr;3(4):513–9.

43. Su Y-C, Yu C-C, Hsu F-T, Fu S-L, Hwang J-J, Hung L-C, et al. Everolimus sensitizes Ras-transformed cells to radiation in vitro through the autophagy pathway. Int J Mol Med. 2014 Nov;34(5):1417–22.

44. Nassim R, Mansure JJ, Chevalier S, Cury F, Kassouf W. Combining mTOR Inhibition with Radiation Improves Antitumor Activity in Bladder Cancer Cells In Vitro and In Vivo: A Novel Strategy for Treatment. PLoS One. 2013;8(6).

45. Pawlik TM, Keyomarsi K. Role of cell cycle in mediating sensitivity to radiotherapy. Int J Radiat Oncol. 2004;59(4):928–42.

46. Wang Z, Lai S-T, Ma N-Y, Deng Y, Liu Y, Wei D-P, et al. Radiosensitization of metformin in pancreatic cancer cells via abrogating the G2 checkpoint and inhibiting DNA damage repair. Cancer Lett. 2015 Aug;

47. Exner S, Prasad V, Wiedenmann B, Grötzinger C. Octreotide does not inhibit proliferation in five neuroendocrine tumor cell lines. Front Endocrinol (Lausanne). 2018;9(APR).

48. Zellmer J, Yen HY, Kaiser L, Mille E, Gildehaus FJ, Böning G, et al. Toxicity of a combined therapy using the mTOR-inhibitor everolimus and PRRT with [177Lu]Lu-DOTA-TATE in Lewis rats. EJNMMI Res. 2020;10(1).

49. Pool SE, Bison S, Koelewijn SJ, van der Graaf LM, Melis M, Krenning EP, et al. mTOR inhibitor RAD001 promotes metastasis in a rat model of pancreatic neuroendocrine cancer. Cancer Res. 2013 Jan;73(1):12–8.

