## Supplemental Fig. 1 and Fig. 2 for "mTOR inhibitors as radiosensitizers in neuroendocrine neoplasms"

### Supplementary Figures

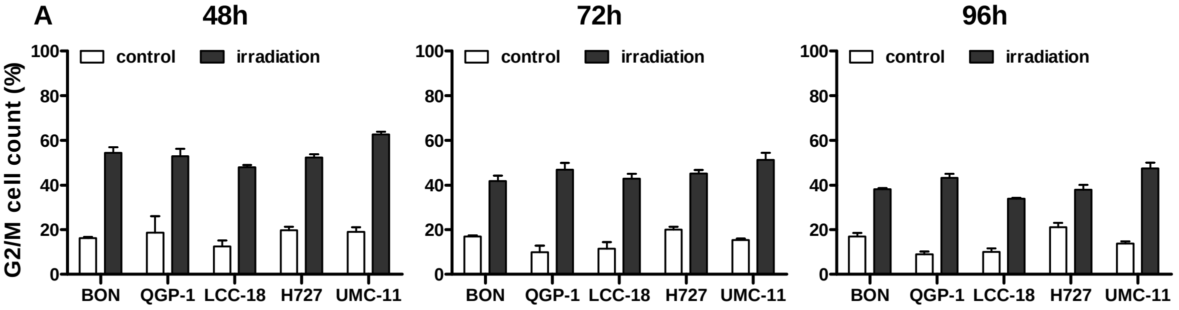


**Figure S1 | Irradiation leads to G2/M accumulation of NET cells after 48, 72 and 96 h.** NET cell lines were irradiated with 10 Gy and samples were collected after the indicated time points for cell cycle analysis by flow cytometer. Data are shown as bar diagrams with mean ± S.E.M. (n=2-3).

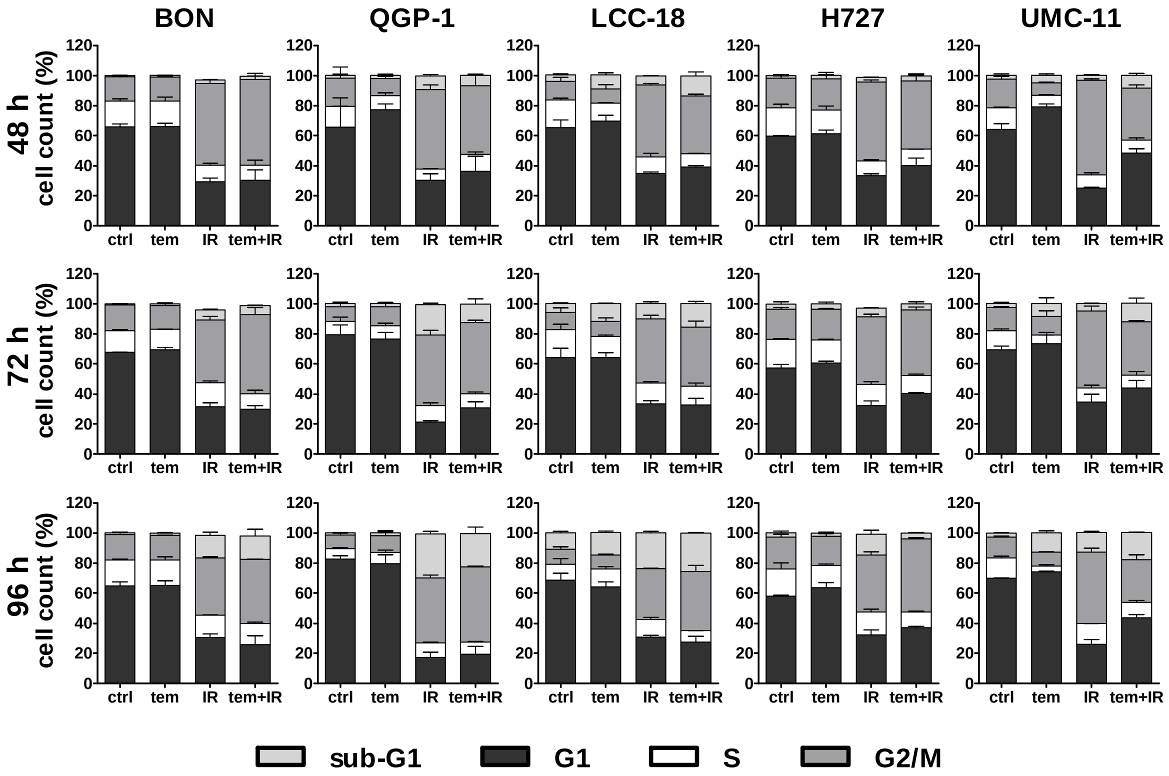


**Figure S2 | Cell cycle distribution at 48, 72 and 96 h after combined treatment.** NET cell lines were incubated with 1 µM temsirolimus (tem) or vehicle (ctrl) for 24 h before irradiation. For assessment of cell cycle distribution pretreated NET cells were collected 48, 72 or 96 h after irradiation with 10 Gy (IR, tem+IR), stained with propidium iodide and analyzed by flow cytometer. Data show mean ± S.E.M. (n=2-3).
